# *Starship* giant transposable elements cluster by host taxonomy using kmer-based phylogenetics

**DOI:** 10.1101/2024.08.30.610507

**Authors:** Rowena Hill, Daniel Smith, Gail Canning, Michelle Grey, Kim E. Hammond-Kosack, Mark McMullan

**Affiliations:** Earlham Institute, Norwich, Norfolk, NR4 7UZ, UK; Intelligent Data Ecosystems, Rothamsted Research, Harpenden, Hertfordshire, AL5 2JQ, UK; Protecting Crops and the Environment, Rothamsted Research, Harpenden, Hertfordshire, AL5 2JQ, UK

**Author notes:** Corresponding authors (R.H.); (K.H-K.); (M.M.). Present address: Department of Computational and Systems Biology, John Innes Centre, Norwich, Norfolk, NR4 7UH, UK.

**Keywords:** cargo-mobilising elements, *Gaeumannomyces*, *Ascomycota*, *Pezizomycotina*

## Abstract

*Starships* are a recently established superfamily of giant cargo-mobilising transposable elements in the fungal subphylum *Pezizomyotina* (phylum *Ascomycota*). To date, *Starship* elements have been identified up to ∼700 Kbp in length and carrying hundreds of accessory genes, which can confer both beneficial and deleterious traits to the host genome. Classification of *Starship* elements has been centred on the tyrosine recombinase gene that mobilises the element, termed the captain. We contribute a new perspective to *Starship* classification by using an alignment-free kmer-based phylogenetic tree building method, which can infer relationships between elements in their entirety, including both active and degraded elements and irrespective of high variability in element length and cargo content. In doing so we found that relationships between entire *Starships* differed from those inferred from captain genes and revealed patterns of element relatedness corresponding to host taxonomy. Using *Starships* from *Gaeumannomyces* species as a case study, we found that kmer-based relationships correspond with similarity of cargo gene content. Our results suggest that *Starship*-mediated horizontal transfer events are frequent between species within the same genus but are less prevalent across larger host evolutionary distances. This novel application of a kmer-based phylogenetics approach overcomes the issue of how to represent and compare highly variable *Starships* elements as a whole, and in effect shifts the perspective from a captain to a cargo-centred concept of *Starship* identity.

**SUMMARY:** We applied a kmer-based phylogenetic classification approach to giant *Starship* cargo-mobilising elements from species across the *Pezizomycotina* (*Ascomycota*, *Fungi*). We found *Starship* elements to frequently cluster according to host taxonomy, suggesting horizontal transfer of elements is less common across larger evolutionary distances. Kmer-based phylogenetics approaches show promise for both element classification and to inform our understanding of the evolution of *Starships* and other giant cargo-mobilising elements.

## INTRODUCTION

Transposable elements (TEs), or transposons, are stretches of DNA which can independently move and replicate within the genome (Biémont 2010; Wells and Feschotte 2020). Thanks to advances in long-read sequencing, highly contiguous genome assemblies have revealed the existence of TEs hundreds of kilobases in length (Arkhipova and Yushenova 2019). Some of these large TEs have been shown to harbour both genes necessary for their mobilisation as well as miscellaneous accessory genes, and are accordingly referred to as cargo-mobilising elements (CME) (Gluck-Thaler and Vogan 2024). Recently, giant CMEs have been found in various species in the fungal subphylum *Pezizomycotina* (phylum *Ascomycota*; McDonald *et al*. 2019; Vogan *et al*. 2021; Urquhart *et al*. 2022), and have since been determined to belong to a newly established TE ‘superfamily’ (*sensu* Wicker *et al*. (2007)) or ‘subclass’ (*sensu* Wells and Feschotte (2020)) known as ‘*Starships’* (Gluck-Thaler *et al*. 2022). To date, *Starship* CMEs have been found to range in length from 15 Kbp (Gluck-Thaler *et al*. 2024) to ∼700 Kbp (Urquhart *et al*. 2023a).

*Starship* mobilisation is mediated by a leading 5’ located gene termed the ‘captain’ containing the DUF3435 domain (protein family accession PF11917), which encodes a tyrosine recombinase that initiates movement of the TE into a new genomic location via a ‘cut-and-paste’ mechanism (Urquhart *et al*. 2023b). This is a similar mobilisation process to the ‘*Crypton’* Class II DNA transposon superfamily (Wells and Feschotte 2020), which was incidentally also first found in fungi (Goodwin *et al*. 2003), although this TE superfamily has since been found in other eukaryotes (Kojima and Jurka 2011). Tyrosine recombinase domains in *Starship* captain genes and *Cryptons* are very distantly related (Gluck-Thaler *et al*. 2022) and, unlike *Cryptons*, *Starship* elements are flanked by direct repeats (DRs) and sometimes tandem inverted repeats (TIRs), and can contain a highly variable and often sizeable cargo of accessory genes (Gluck-Thaler and Vogan 2024). *Starship* cargos can harbour genes that are beneficial to the fungus, for example those associated with plant virulence (McDonald *et al*. 2019), metal tolerance (Urquhart *et al*. 2022) and climate adaptation (Tralamazza *et al*. 2024). However, as selfish genetic elements, *Starship* may also mobilise cargo which is neutral or even detrimental to the overall fitness of the host genome (Vogan *et al*. 2021).

Until now, classification within the *Starship* CME superfamily has been focused on the captain gene, using both phylogenetic relationships between captain and other tyrosine recombinase genes to define ‘family’ and orthologue clustering of captain genes to define ‘navis’ (i.e. a ship) (Gluck-Thaler and Vogan 2024). *Starship* captain genes do not form a single monophyletic cluster in the tyrosine recombinase gene tree, and are instead scattered across the phylogeny amongst other tyrosine recombinase genes (Gluck-Thaler *et al*. 2022; Hill *et al*. 2024). Due to their highly divergent nature, tyrosine recombinase gene sequences are also difficult to align, introducing uncertainty into conventional alignment-based phylogenetic analyses. It is not currently possible to determine whether these relationships described by captains are preserved or representative of the *Starships* as a whole, considering that elements are highly variable in terms of cargo and overall length. This also limits assessment of the prevalence of (or boundaries to) horizontal exchange across the *Pezizomycotina*. In an effort to represent distinction in cargo content, Gluck-Thaler and Vogan (2024) introduced the additional definition of ‘haplotype’, based on clustering of kmer similarity scores. Here, we have taken this approach one step further and used a kmer-based phylogenetic tree building method to contribute a new perspective to *Starship* classification. In doing so we have revealed previously obscured patterns of *Starship* relatedness corresponding to host taxonomy. *Starship* horizontal transfer events remain apparent but are less prevalent across large evolutionary distances.

To determine whether the relatedness revealed by the kmer trees conformed with similarity in cargo gene content, we explored the cargos of *Starships* previously identified from genomes within the genus *Gaeumannomyces* (Hill *et al*. 2024), which comprises both pathogenic and non-pathogenic wheat and wild grass associates (Palma-Guerrero *et al*. 2021; Chancellor *et al*. 2024). These elements provided an ideal case-study as they vary greatly in overall size and number of cargo genes within their host-taxonomy clusters. The genomes were also all generated in parallel using the same long-read sequencing technology and a cross-referent annotation pipeline (Hill *et al*. 2024). Given the impact of assembly and annotation quality on *Starship* recovery (Gluck-Thaler and Vogan 2024), these *Gaeumannomyces* elements therefore represent a consistent dataset that are impacted to a lesser extent by the technology used to produce them.

## MATERIALS AND METHODS

### Kmer-based phylogenetic classification

To compare phylogenetic reconstruction of whole elements versus captain genes, we used a curated set of 39 *Starships* from Gluck-Thaler *et al*. (2022) and Gluck-Thaler and Vogan (2024) alongside 20 *Gaeumannomyces Starships* predicted using the tool starfish v1.0.0 (Gluck-Thaler and Vogan 2024) in our previous study (Hill *et al*. 2024). We used entire element sequences as input for the kmer-based method Mashtree v1.4.6 (Katz *et al*. 2019) with 1,000 bootstrap replicates and the --min-depth 0 parameter to discard very unique kmers, recommended to improve accuracy. We used the corresponding captain genes as input for a maximum likelihood (ML) tree, first aligning gene sequences using MAFFT v7.271 (Katoh and Standley 2013), trimming using trimAl v1.4.rev15 (Capella-Gutiérrez *et al*. 2009), and finally building the ML tree using RAxML-NG v1.1.0 (Kozlov *et al*. 2019) with bootstrapping until convergence, which occurred after 150 bootstrap replicates. We visualised concordance between the two phylogenies via a tanglegram, produced in R v4.3.1 (R Core Team 2022) using the packages ape v5.7-1 (Paradis and Schliep 2019), phytools v2.1-1 (Revell 2024) and ggtree v3.9.1 (Yu *et al*. 2017). We calculated the normalised Robinson–Foulds (RF) distance between the element and captain phylogenies using the RF.dist function from the phangorn v2.7.0 package (Schliep *et al*. 2017).

We then used a larger dataset of *Starships* predicted using the tool starfish v1.0.0 by Gluck-Thaler and Vogan (2024) to assess whether patterns in the curated kmer tree would persist with broader sampling. In total, 617 entire element sequences were again run with Mashtree, but with 100 bootstrap replicates and the default –min-depth parameter to accommodate for the much larger dataset. Previously determined *Starship* family classification based on captain phylogenetic relationships (Gluck-Thaler and Vogan 2024) were mapped to tree tips to visualise the distribution of families across clades using the additional R packages ggtreeExtra v1.10.0 (Xu *et al*. 2021) and glottoTrees v0.1.10 (Round 2021).

### Exploration of cargo gene content in *Gaeumannomyces* elements

We used the aforementioned dataset of twenty *Starships* predicted from seven *Gaeumannomyces* genomes to assess whether similarities in cargo gene content corresponded with the patterns of relatedness described by the kmer trees. We characterised orthologous genes predicted in our previous study (Hill *et al*. 2024) as being core, accessory or specific within the set of twenty elements, and their sharedness was visualised using the R package ComplexUpset v1.3.3 (Krassowski 2022). After normalising cargo orthogroup presence-absence values with the base R scale function, we produced a Euclidean distance matrix using the R dist function and performed hierarchical clustering with the hclust function using the ‘complete’ agglomeration method. We then compared the topology produced by hierarchical clustering with phylogenetic relationships from the larger kmer-based tree using a tanglegram and calculated the normalised RF distance, as described above. We also determined the location of cargo orthogroups – i.e. whether orthologous genes were only found inside elements or also found in the wider genome.

We searched for specific genes or domains previously reported to be prevalent in *Starships* or of genes with assigned functional roles of particular note (Gluck-Thaler *et al*. 2022) using BLAST v2.10 (Camacho *et al*. 2009) and also PFAM domain assignment from the functional annotation (Hill *et al*. 2024). Namely: DUF3723, ferric reductase (FRE), patatin-like phosphatase (PLP), ToxA effector, spore killing (Spok) genes 1,2,3 and 4, and associated domains. We additionally made BLAST searches against the Pathogen–Host Interactions Database v4.17 (PHI-base; Urban *et al*. 2022) downloaded on 1^st^ August 2024, and considered a positive match when at least 50% of genes in an orthogroup had the same hit. We assessed whether gene ontology (GO) terms were enriched amongst cargo genes using the R package topGO v2.52.0 (Alexa and Rahnenfuhrer 2022) with Fisher’s exact test and the weight01 algorithm.

In addition to previously mentioned packages, data analysis and visualisation was done using the following R packages: cowplot v1.1.3 (Wilke 2024), ggforce v0.4.2 (Pedersen 2024), gggenomes v1.0.0 (Hackl *et al*. 2024), ggnewscale v0.4.10 (Campitelli 2024), ggpubr v0.6.0 (Kassambara 2023), ggrepel v0.9.5 (Slowikowski 2024), matrixStats v1.3.0 (Bengtsson 2024), patchwork v1.2.0 (Pederson 2024), scales v1.3.0 (Wickham and Seidel 2023), tgutil v0.1.15 (Chomsky and Lifshitz 2023) and tidyverse v2.0.0 (Wickham *et al*. 2019).

## RESULTS AND DISCUSSION

### A kmer-based approach to *Starship* phylogenetic classification recovers signal corresponding to host taxonomy

We used a novel kmer-based approach for phylogenetic classification of *Starships* to produce a phylogenetic tree from 59 entire *Starship* element sequences, encompassing 17 host genera across 6 classes in the *Pezizomycotina*. We found elements to broadly cluster by genus, even when differing greatly in length (Fig. 1a). This contrasted with the captain gene tree (Supplementary Fig. 1) and element and captain trees were frequently discordant (RF distance 0.74=74% differing bipartitions; Fig. 1b), i.e. *Starships* that were more closely related according to their kmer profiles could have very divergent captain genes. There were some exceptions to element/captain discordance, for instance conserved relationships in both captain and element trees were observed for the *Alternaria* clade (Fig. 1b). The degree of element/captain phylogenetic discordance is important not only because phylogenetic relationships of captains have been the predominant factor in element classification (Gluck-Thaler and Vogan 2024), but also because similar captain genes being found in host genomes from distant clades of the fungal tree of life may have been taken as evidence for *Starship* transfer between distantly related species. Also note the placement of Mpha_Derelict – a previously ‘unclassifiable’ deactivated *Starship* missing the captain gene – alongside other elements from *Macrophomina* species (Fig. 1a).

**Fig. 1.**
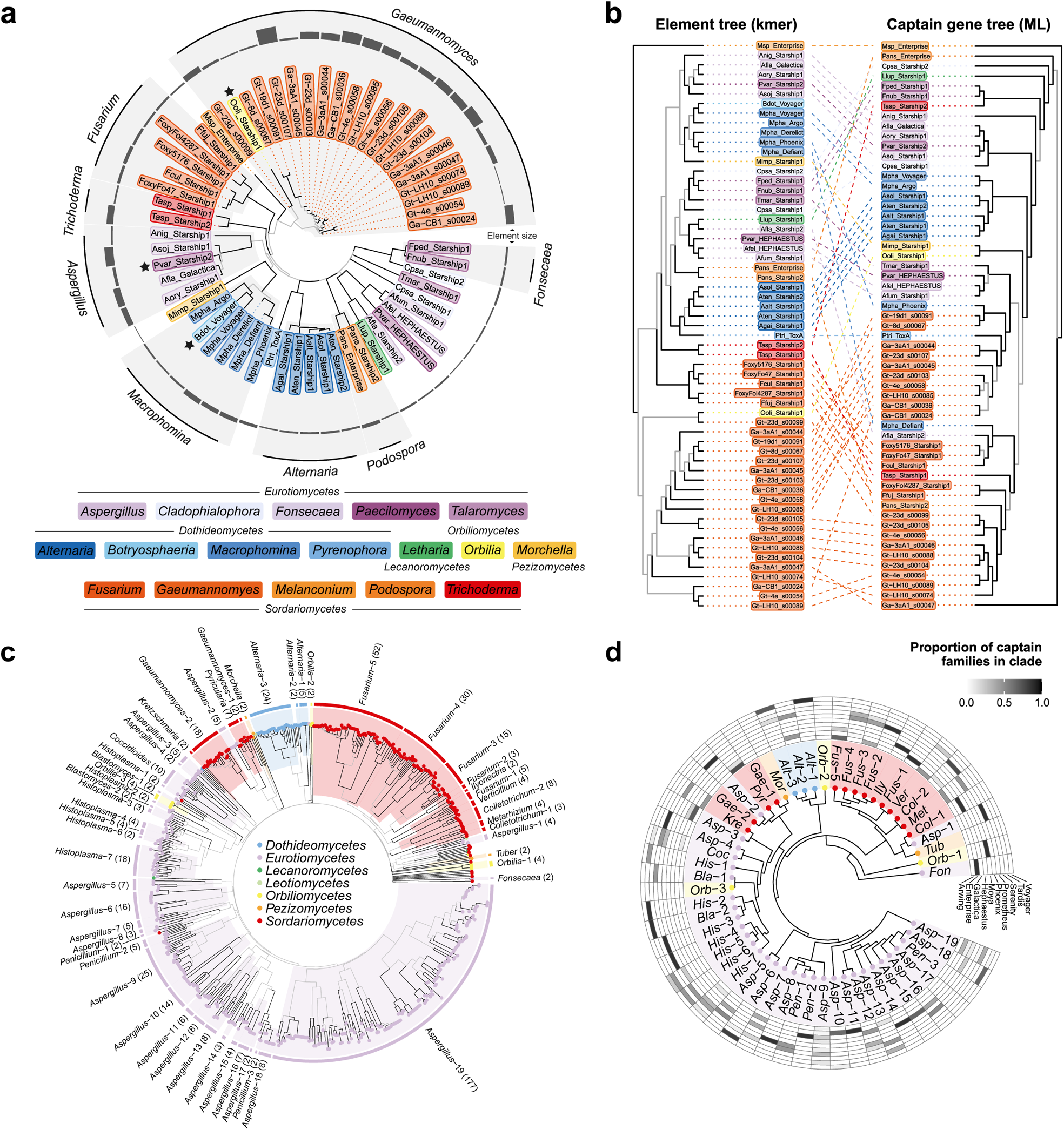
Kmer-based phylogenetic analyses of *Starship* elements. a) An unrooted kmer-based phylogenetic tree of 59 *Starships* – 39 curated elements from 33 *Pezizomycotina* species (Gluck-Thaler *et al*. 2022; Gluck-Thaler and Vogan 2024) and 20 predicted by starfish from *Gaeumannomyces* species (Hill *et al*. 2024). Grey branches indicate bootstrap support < 70. Tip points are coloured by genus and the outer ring indicates total element length. Black stars beside tips highlight elements from another genus in an otherwise monophyletic clade. b) A tanglegram comparing the topology of the kmer-based element tree in Fig. 1a and a maximum-likelihood gene tree of the corresponding captain genes (see Supplementary Fig. 1 for the unrooted captain tree). Both trees are arbitrarily rooted with the Msp_Enterprise element. Grey branches indicate bootstrap support < 70. c) An unrooted kmer-based phylogenetic tree of 617 *Starships* predicted with starfish (Hill *et al*. 2024; Gluck-Thaler and Vogan 2024), with grey branches indicating bootstrap support < 70. Genus-level monophyletic clades are highlighted and labelled, with the number of elements in each clade shown in brackets. Clades and tips are coloured by host taxonomic class. d) A summary of the kmer-based tree in Fig. 1c with genus-level monophyletic clades collapsed. The outer grid summarises *Starship* family classifications based on captain genes for the elements in each clade, with a darker grid cell colour indicating a higher proportion of the elements within the clade belonging to that family.

To see if these observations of clustering by host taxonomy extended more broadly across the *Pezizomycotina*, we used the same kmer-based phylogenetics method on a larger dataset of 617 elements systematically predicted using the tool starfish (Gluck-Thaler *et al*. 2024). This again recovered widescale clustering by host taxonomy, with the additional clear formation of clades broadly corresponding to host class-level (Fig. 1c). Our results suggest that *Starships* more readily horizontally transfer amongst closely related species and are less likely to transfer between species separated by greater evolutionary distances. Importantly, there is evidence for some exceptional major horizontal transfer events, as monophyletic clades are occasionally interspersed with elements from more distantly related hosts, for example see the *Aspergillus*-2 clade (Supplementary Fig. 2). However, the evolutionary distances across which *Starships* can be, or are likely to be, exchanged may have been inflated from results based only on phylogenetics of captain genes.

As may be expected from the observed element/captain tree discordance, family classifications based on captains were scattered across the larger starfish-predicted element kmer tree (Fig. 1d). There were many within-genus subclades of elements with captains of the same family, but also cases where minimally diverged sister elements had different captains – for example, aspcri2_s00912/aspcri1_s00891 (Phoenix/Prometheus, respectively) in different genomes, and aspnig6_s01954/aspnig6_s01955 (Hephaestus/Phoenix, respectively) in the same genome (Supplementary Fig. 2). A similar observation was made by Gluck-Thaler and Vogan (2024) for *Starship* pairs with near identical cargo ‘haplotypes’ but different captain-derived families. Together with the fact that captain genes are phylogenetically indistinguishable from ‘lone’ tyrosine recombinase genes harbouring the DUF3435 domain (Gluck-Thaler *et al*. 2022; Hill *et al*. 2024), this prompts the question as to whether *Starships* can swap the captain for a different tyrosine recombinase gene, which would render the ‘captain’ status as somewhat transient. A previous study has already reported that *Starship* elements can lose their captain gene to become ‘degraded’ or ‘derelict’ (Gluck-Thaler *et al*. 2022), and in another study a mechanism has been suggested wherein captain inactivation enables cargo swapping (Urquhart *et al*.2023a). A similar mechanism where the captain, rather than the cargo, is swapped to acquire a captain gene from a different family could be a strategy to diversify insertions of virtually identical elements into different target sites.

Aside from the major clade in the larger starfish kmer tree overrepresented with elements from eurotiomycete hosts (Fig. 1c), other eurotiomycete elements appeared scattered amongst other clades, with a conspicuous *Aspergillus-Trichophyton* clade dividing sordariomycete elements from *Gaeumannomyces*, *Pyricularia* and *Sporothrix* species (Supplementary Fig. 2). It is notable that eurotiomycete elements dominate the starfish dataset – of all the genomes explored by Gluck-Thaler and Vogan (2024), *Eurotiomycetes* was the class with the highest proportion of genomes returning a *Starship* (36%; Supplementary Fig. 3). This was closely followed by the *Orbiliomycetes* (28%), despite 16 times fewer orbiliomycete genomes having been surveyed compared to the *Eurotiomycetes*, and orbiliomycete element clades were similarly widespread across the kmer tree (Fig. 1c). As one of the earliest diverging classes within the *Pezizomycotina* subphylum, the *Orbiliomycetes* are distantly related to *Eurotiomycetes* (Li *et al*. 2021), and they do not particularly share ecological distributions more so than other taxonomic classes, so the underlying biological explanation is unclear. The far larger *Eurotiomycetes* class comprises diverse lifestyles including rock-inhabiting fungi and other extremophiles; plant and animal pathogens; lichenised and lichen-associated fungi; ectomycorrhizal fungi; ant mutualists; and saprotrophs (Geiser *et al*. 2015). The *Orbiliomycetes* are primarily thought to be saprotrophs but include some soil dwelling carnivorous fungi which trap invertebrates (Pfister 2015). Variation in the rate of *Starship* recovery in the genomes of different taxonomic classes could be a result of inconsistencies in assembly quality, or bias within the starfish tool to recover certain elements from certain classes. However, these results do suggest that there may be a relationship between the tendency for a taxonomic class to have *Starship* elements and the host evolutionary distance these CMEs may transfer across.

While we consider this to be a promising application for kmer-based phylogenetics methods, we must note that such methods were typically developed for whole-genome input. We are not aware of kmer-based phylogenetic methods having been tested on sequences such as fungal CMEs. However, given that such methods are considered well suited to viral genomes due to their high levels of mutation, gene duplication and rearrangement (Zielezinski *et al*. 2017), CMEs would appear to be a similarly appropriate use-case. There are also many different approaches and tools for alignment-free sequence comparison which would warrant further testing in this context (Luczak *et al*. 2019; Zielezinski *et al*. 2019). We were unable to produce a kmer tree for captain genes using Mashtree, presumably due to the much smaller sequence length of a single gene. This meant we were not able to directly compare whole element and captain trees using the same kmer-based method. However, at the genome-scale, previous comparisons of alignment and kmer methods suggest reasonable topological congruence (Van Etten *et al*. 2023), or no greater incongruence than might be expected from using different alignment-based methods (e.g. Shen *et al*. 2021).

There are some limitations to alignment-free phylogenetics methods. Unlike conventional alignment-based phylogenetic trees, alignment-free trees do not produce branch lengths with a scale corresponding to geological time, and so one cannot extrapolate the date of divergences. Alignment-free methods also struggle with the reconstruction of deep nodes (Fan *et al*. 2015), which is evident from the kmer trees we present here. This may limit the ability of these methods to address questions about inter-relatedness of larger CME clades but should still allow for the detection of more recent horizontal transfer events.

### Both cargo genes and non-coding cargo content contribute to kmer-based phylogenetic relationships between *Gaeumannomyces Starships*

In order to explore the extent to which cargo gene content corresponded with the kmer-based phylogenetic relationships, we used twenty *Starships* previously identified from seven genomes across three separate lineages within the genus *Gaeumannomyces,* an understudied member of the *Magnaporthaceae* (Hill *et al*. 2024). These genomes were sequenced from five strains of the wheat root pathogen species *G. tritici* (*Gt*) and two of the oat root pathogen *G. avenae* (*Ga*). Within the *Gt* strains there is further subdivision of two strains belonging to ‘type A’ and three to ‘type B’, two distinct genetic lineages present in the species (Palma-Guerrero *et al*. 2021). This division is meaningful, as differences between the two types in terms of both virulence and genomic signatures may indicate that these two types actually represent cryptic species (Hill *et al*. 2024). As well as being a consistently amassed set of *Starships* for controlled comparison, these *Gaeumannomyces* elements also provided major variability, ranging from ∼32–688 Kbp in total length and containing between 1–156 genes.

We found that *Starships* with greater numbers of shared orthologous genes were frequently sister elements or closely related in the kmer tree, for instance Gt-LH10_s00088, Gt-23d_s00104 and Ga-3aA1_s00046 (Fig. 2a). Most cases of more distantly related elements with high cargo gene sharedness involved the largest and most gene-rich element, Gt-23d_s00107, which incidentally also had one of the highest proportions (48%) of element-specific genes. Hierarchical clustering of cargo orthologous gene content supported these results, with reasonable concordance between the hierarchical clustering and kmer element tree (RF distance 0.47=47% differing bipartitions; Fig. 2c) and the most notable deviation between the two trees was the divergence of element Gt-23d_s00107. While cargo gene content was evidently a contributing factor to the patterns of *Gaeumannomyces* element relatedness recovered in the kmer-based phylogenies, the nature of a kmer-based approach means that intergenic content within *Starships* must also be implicated. Indeed, repetitive DNA, introns and presumably other non-coding regions can provide important phylogenetic signals (Lo *et al*. 2022). Here, the only two *Gt*A elements found, one in each *Gt*A genome, contained a single cargo gene despite being 61 and 73 Kbp long. These elements were nonetheless found to cluster with other *Gaeumannomyces* species in the curated *Starship* tree (Fig. 1a). Wider sampling of starfish-predicted elements in the larger kmer tree revealed more nuance, as the *Gt*B and *Ga* elements were closely related to the rice blast fungus *Pyricularia oryzae* (syn. *Magnaporthe oryzae*) elements, while the single-gene *Gt*A elements, were in a distinct clade more closely related to an element from the *Sordariomycetes* species *Sporothrix brasiliensis*, albeit without significant branch support (Supplementary Fig. 2). *S. brasiliensis* is found in soils and vegetation, but is also an opportunistic mammalian pathogen, primarily of humans and cats, due to its temperature-dependent dimorphic lifestyle (Téllez *et al*. 2014). Despite being a similar length (57 Kbp) to the *Gt*A elements, the *S. brasiliensis* element contained 19 genes, none of which showed sequence similarity with the single gene found in the *Gt*A elements. This suggests that it was primarily non-coding cargo content that informed kmer-based relationships between the *S. brasiliensis* and *Gt*A elements.

**Fig. 2.**
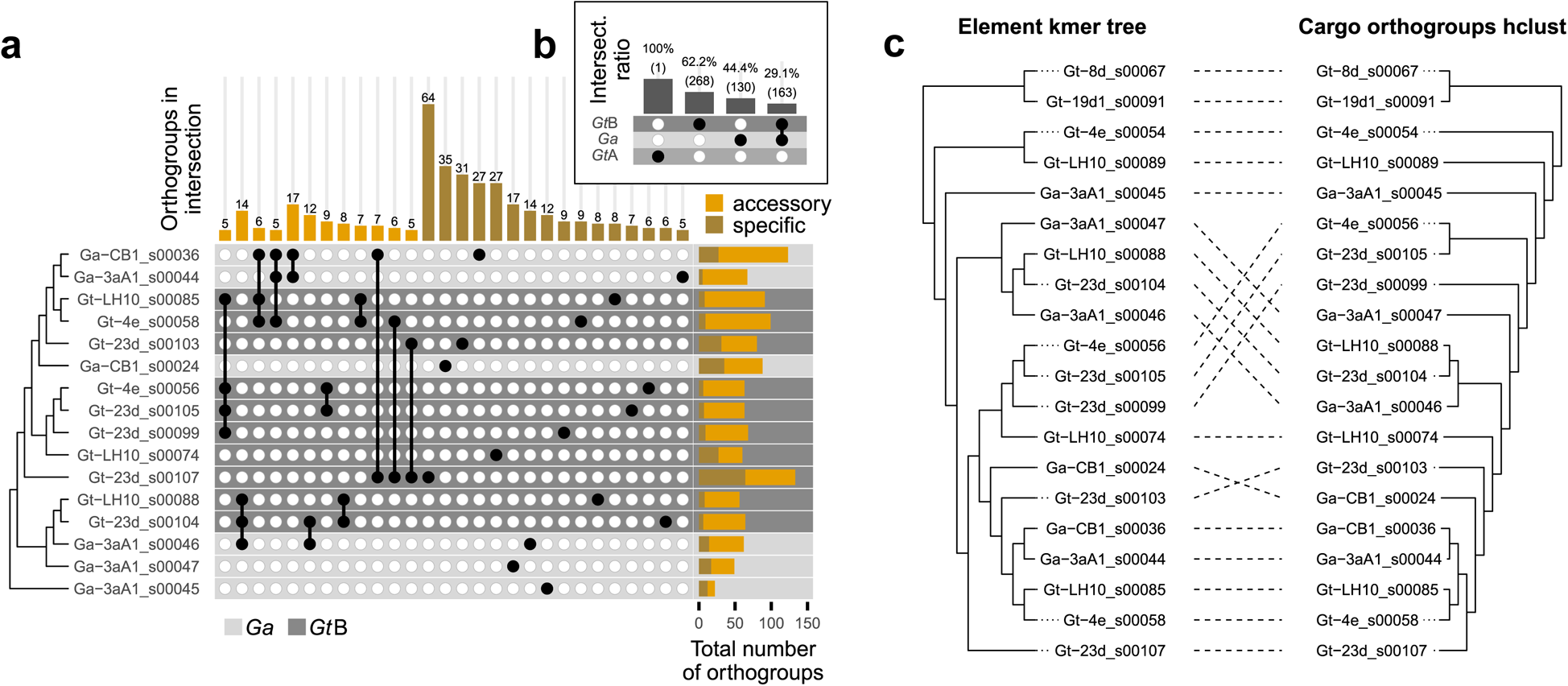
Comparison of cargo gene content similarity across *Gaeumannomyces Starships*. a) An upset plot indicating groups of elements which share at least five orthologous genes (accessory), and elements with at least five unique cargo genes (specific). Elements are ordered by phylogenetic relationships taken from Supplementary Fig. 2. Total number of cargo orthogroups is shown in the right-hand bar plot with the proportion of accessory and specific cargo genes coloured per element. Element rows are coloured by host lineage. See a representation of all shared accessory orthologous genes in Supplementary Fig. 4. (b) An upset plot indicating the ratio of orthologous genes shared across lineage/species boundaries. (c) A tanglegram comparing the phylogenetic relationships of *Gaeumannomyces* elements taken from Supplementary Fig. 2 and a hierarchical clustering of cargo orthologous gene presence/absence.

Both the kmer trees and the patterns of shared cargo genes indicated that there is no apparent species boundary for *Starship* exchange between *Gt*B and *Ga.* We found no evidence of exchange with *Gt*A strains, although there was only one gene-poor *Gt*A element with which to compare. Nonetheless, together with the fact that, unlike the other *Gaeumannomyces* elements, the *Gt*A elements were previously found to be subject to element-wide repeat induced point mutation (RIP) (Hill *et al*. 2024), *Starship* prevalence and exchange limits may be another symptom of cryptic speciation between *Gt* types. Although *Gt*B and *Ga* elements appear to be readily exchanged, there was an imbalance in how cargo genes were shared, as a higher proportion of *Ga* cargo genes had an orthologue in *Gt*B elements (56%) than *Gt*B cargo genes had in *Ga* elements (38%; Fig. 2b). Additionally, there were differences in how cargo genes were distributed in the genome, with more cargo gene orthogroups only found inside *Ga* elements that had copies integrated into the wider genome in *Gt* strains than the reverse (Supplementary Fig. 5a). In a similar vein, *Ga Starships* broadly had a higher proportion of orthogroups that were only inside the element compared to *Gt*B *Starships* (Supplementary Fig. 5b). Unpicking the differences in relative levels of duplication, sharedness and location of cargo genes on different *Starships* may be important for determining patterns of directionality or inheritance in element exchange.

### *Gaeumannomyces Starship* cargos harbour a variety of putative plant–fungal interaction genes, but the ToxA gene was notably absent

Most genes previously reported to be common in *Starships* (Gluck-Thaler *et al*. 2022) were notably absent from *Gaeumannomyces Starship* cargos, namely DUF3723, ferric reductase (FRE), patatin-like phosphatase (PLP) and spore-killer (Spok 1,2,3 and 4). There was one putative nucleotide-binding domain and leucine-rich repeat containing gene (NLR) located on element Gt-23d_s00107 (Fig. 3). The NLR contained a central NACHT domain – the most common nucleotide binding and oligomerization (NOD) domain in fungal NLRs (Daskalov *et al*. 2020) – a WD40 repeat domain, and a sesA N-terminal domain of unknown function (PF17107) that is more common in ascomycete NLRs (Daskalov *et al*. 2020). This sesA-NACHT-WD structure is also found in the NWD3 gene of *Podospora anserina* (Daskalov *et al*. 2012). While the function of sesA is not established, other members of the *P. anserina* NWD gene family are involved in heterokaryon/vegetative incompatibility or self/non-self recognition, which has also been hypothesised to contribute to an innate fungal immune system (Paoletti and Saupe 2009; Uehling *et al*. 2017). Of particular note was the absence of ToxA in the *Gaeumannomyces* cargos, which is a necrotrophic effector horizontally transferred by *Starships* in three other wheat pathogens – *Pyrenophora tritici-repentis, Parastagonospora nodorum* and *Bipolaris sorokiniana* (McDonald *et al*. 2019; Bucknell *et al*. 2024). It is highly likely that *Gaeumannomyces* spp. cooccur with one or more of these wheat pathogens, which would have provided the opportunity to exchange *Starships.* However, all three species containing ToxA reside in a different class, *Dothideomycetes*, in the order *Pleosporales*. At the present time, the lack of ToxA in the *Gaeumanomyces Starships* is consistent with our kmer tree results indicating a host relatedness boundary to *Starship* exchange.

**Fig. 3.**
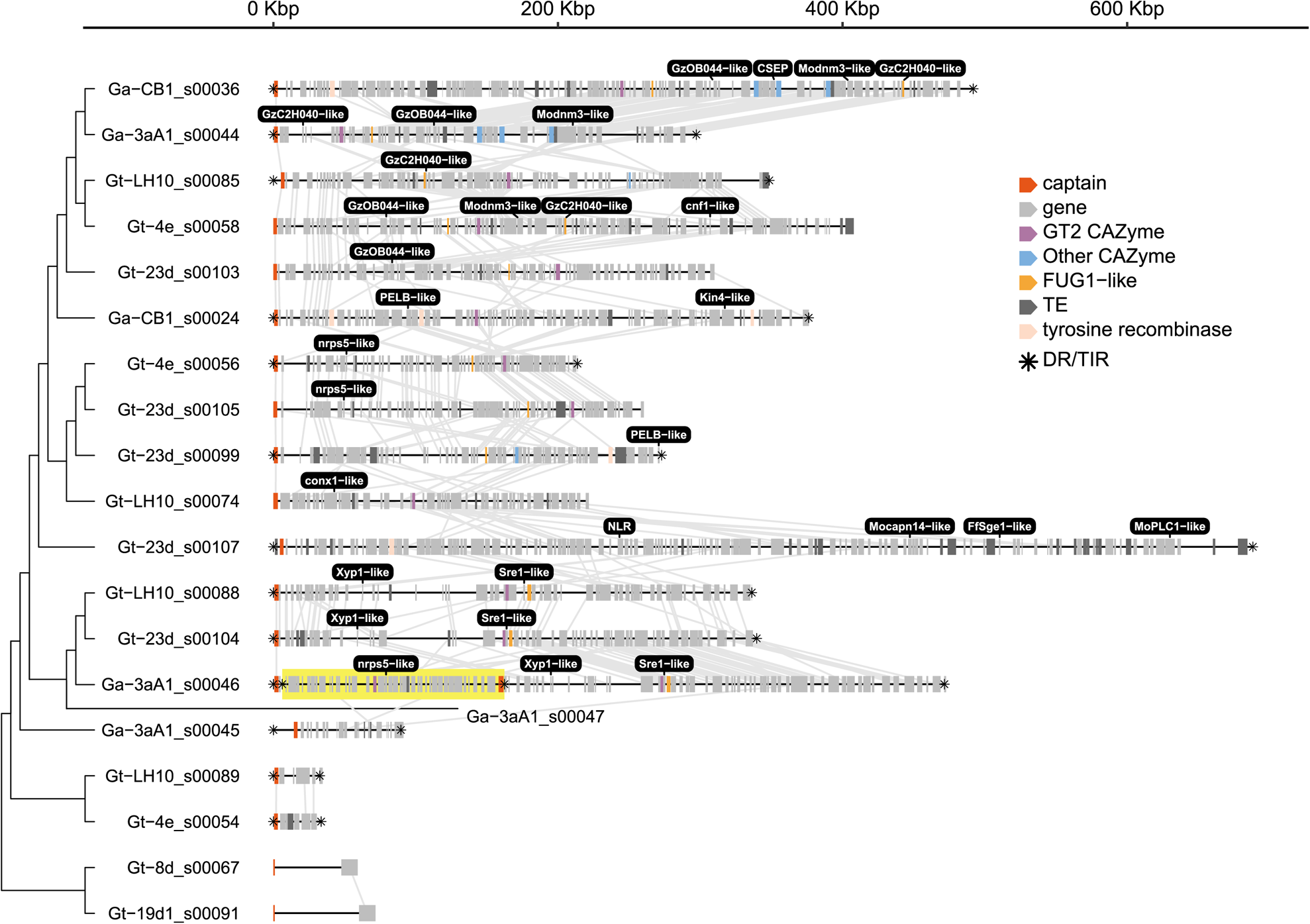
A schematic of the 20 *Starships* predicted using starfish by Hill *et al*. (2024), ordered by phylogenetic relationships taken from Supplementary Fig. 2. Synteny between orthologous genes in neighbouring elements is indicated with grey lines. A nested element is highlighted in yellow. Common genes are coloured with known functions and the presence of flanking direct repeats (DRs) or tandem inverted repeats (TIRs) are indicated with an asterisk. Genes of note are labelled in black boxes.

Regarding whether the *Gaeumannomyces Starship* cargos exhibited a core functional role, GO term enrichment analysis of cargo genes reflected high variability as there was no significant enrichment in most elements, although ubiquinone biosynthesis and regulation of translational fidelity were significantly enriched in Ga-3aA1_s00044 and Ga-CB1_s00036, respectively. There were no cargo orthogroups that were core to all elements, but five orthogroups were present in at least 50% of the elements (Supplementary Fig. 4). One was predicted to be a carbohydrate-active enzyme (CAZyme) belonging to glycosyltransferase family 2 (GT2; Fig. 3). The GT2 family includes enzymes necessary for the synthesis of chitin (Lairson *et al*. 2008), which is required for structural integrity of the fungal cell wall (Bowman and Free 2006). A GT2 enzyme has been demonstrated to be required for the disease-causing abilities of the wheat pathogens *Zymoseptoria tritici* and *Fusarium graminearum* (King *et al*. 2017). Expansion and contraction of GT2 CAZyme genes have been shown to be strong predictors of phytopathogenicity and saprotrophy respectively (Dort *et al*. 2023), but GT2 genes are also expanded in mycorrhizal lineages (Rosling *et al*. 2024), suggesting a key role in both pathogenic and mutualistic plant– fungal interactions. In addition to the prevalent GT2 orthogroup, other CAZymes and CAZyme families were found in various elements: sterol 3β-glucosyltransferase (GT1), glycoside hydrolase (GH) family 33, α-galactosidase (CBM35+GH27), and glucose-methanol-choline oxidoreductase (AA3_2) in elements Ga-3aA1_s00044 and Ga-CB1_s00036; chitinase (GH18) in Gt-LH10_s00085; and another GT2 CAZyme in Gt-23d_s00099.

Multiple *Gaeumannomyces Starship* cargo genes had BLAST hits to genes in the PHI-base database, which compiles and curates experimentally verified genes implicated in pathogen– host interactions (Urban *et al*. 2022). This included four genes in closely related *P. oryzae* which have been associated with virulence in barley and rice, two of which are implicated in calcium signalling and two transcription factors, and the previously mentioned GT2 CAZyme which has been associated with virulence of *Zymoseptoria tritici* and *Fusarium graminearum* in wheat leaves and floral spikes, respectively (Table 1). Intriguingly, the chitinase CAZyme cargo gene in element Gt-LH10_s00085 matched a chitinase gene in the mycoparasite *Trichoderma virens* which is associated with its virulence towards the basidiomycete plant pathogen *Rhizoctonia solani*. *Trichoderma* species are known for endophytic colonisation of plants, particularly roots, and in some cases can reduce disease via both inducing plant resistance and direct antagonism of other fungi (Harman *et al*. 2004). Two further orthogroups had hits to CAZyme genes in PHI-base (Xyp1 and PELB/CcpelA), however as these were not previously flagged during CAZyme annotation (Hill *et al*. 2024) there remains some uncertainty as to their function.

**Table 1.** PHI-base genes with BLAST hits in *Gaeumannomyces Starship* cargos. * Pectate lyases CcpelA and PELB matched to the same orthogroup.

| PHI-base ID | Gene | Function | Species | Mutant phenotype | Plant host |
| --- | --- | --- | --- | --- | --- |
| PHI:7559 | FgGT2 | glycosyltransferase | *Fusarium graminearum* | loss of pathogenicity | *Triticum aestivum* |
| PHI:2057 | MoPLC1 | modulator of calcium flux | *Pyricularia oryzae* † | loss of pathogenicity | *Oryza sativa* |
| PHI:3837 | Sre1 | iron-sensitive transcription factor | *Bipolaris maydis* | reduced virulence | *Zea mays* |
| PHI:2476 | CcpelA * | pectate lyase | *Colletotrichum coccodes* | reduced virulence | *Solanum lycopersicum* |
| PHI:222 | PELB * | pectate lyase | *Colletotrichum gloeosporioides* | reduced virulence | *Persea americana* |
| PHI:9042 | nrps5 (FGSG_13878) | non-ribosomal peptide synthetase | *Fusarium graminearum* | reduced virulence | *Triticum aestivum* |
| PHI:6262 | FUG1 | role in pathogenicity and fumonisin biosynthesis | *Fusarium verticillioides* | reduced virulence | *Zea mays* |
| PHI:3315 | conx1 | Zn<sup>2</sup>Cys<sup>6</sup> transcription factors | *Pyricularia oryzae* † | reduced virulence | *Oryza sativa* |
| PHI:3308 | cnf1 | Zn<sup>2</sup>Cys<sup>6</sup> transcription factors | *Pyricularia oryzae* † | reduced virulence | *Hordeum vulgare* |
| PHI:2113 | Kin4 | Ca<sup>2+</sup>/CAM-dependent serine/threonine protein kinases | *Pyricularia oryzae* † | reduced virulence | *Hordeum vulgare* |
| PHI:144 | CHT42 | chitinase | *Trichoderma virens* | reduced virulence | *Rhizoctonia solani* |
| PHI:3210 | FfSge1 | morphological switch regulator | *Fusarium fujikuroi* | unaffected pathogenicity | *Oryza sativa* |
| PHI:1603 | GzOB044 | transcription factor | *Fusarium graminearum* | unaffected pathogenicity | *Triticum* |
| PHI:1377 | GzC2H040 | transcription factor | *Fusarium graminearum* | unaffected pathogenicity | *Triticum* |
| PHI:6639 | Modnm3 | dynamain | *Pyricularia oryzae* † | unaffected pathogenicity | *Oryza sativa* |
| PHI:6613 | Mocapn14 | calpain | *Pyricularia oryzae* † | unaffected pathogenicity | *Oryza sativa* |
| PHI:124206 | Xyp1 (Uv8b_02447) | cell wall degrading enzymes | *Ustilaginoidea virens* | increased virulence | *Oryza sativa* |

Also of note is that none of the biosynthetic gene clusters (BGCs) previously identified in the *Gaeumannomyces* genomes were present in any *Starships*, but two cargo genes had hits to PHI-base genes implicated in secondary metabolite synthesis in *Fusarium* species: nrps5 and FUG1. The latter is involved in fumonisin synthesis in *Fusarium verticillioides* (Ridenour and Bluhm 2017), but is located on a separate locus to the fumonisin (FUM) gene cluster, suggesting that it may play a regulatory role, as biosynthesis transcription factors can frequently be located outside of contiguous BGCs (Kwon *et al*. 2021). FUG1 was also previously found to have orthologues across *Ascomycota*, including in *Gt* (Ridenour and Bluhm 2017). The non-ribosomal peptide synthetase nrps5 gene is located alongside nrps9 in an eight-member BGC cluster in *Fusarium* species, which produces fusaoctaxin A and is essential to virulence of *F. graminearum* in wheat (Jia *et al*. 2019). However, none of the genes surrounding the nrps5-like gene in the *Gaeumannomyces* elements showed similarity to the other nrps5/9 cluster members. We also found an uncharacterised candidate secreted effector protein (CSEP) gene in one element (Ga-CB1_s00036). Intriguingly this CSEP was located within a region that was highly syntenic with another element (Ga-3aA1_s00044) but the CSEP was not present in that second element (Fig. 3), underlining the dynamism of *Starship* cargos.

## Conclusions

Here, we provide evidence of a difference in evolutionary history between *Starship* elements in their entirety versus their captain genes. This raises the question: is it more important to define *Starships* by their mode of mobilisation – i.e. the tyrosine recombinase captain gene – or the cargo of genes and non-coding/repetitive content mobilised? The answer to that question will depend on the context in which the question is asked; namely, whether the enquiry at hand is to understand the mechanism of transposition, or to understand how elements evolve and impact host fitness. Whole-element relationships are easily assessed using kmer-based phylogenetic methods, which has revealed previously hidden signals corresponding to host taxonomy. These methods also allow us to assess relationships including ‘degraded’ elements and detect potential horizontal transfer events where captain and/or DRs/TIRs have been lost. By accounting for the composition of *Starships* without being hampered by alignment issues caused by repeats, indels, duplications, rearrangements and inversions, or lack of available sequences in general, kmer-based phylogenetic methods can help to refine classification of CMEs. Beyond informing classification, this new approach could also provide context and new insights to address fundamental outstanding questions regarding *Starships* and other CMEs, such as the evolutionary origins of elements, the basis for element exchange limits and the role of elements in the host genome.

## DATA AVAILABILITY

All original data sources used in this study are cited in the text. Analysis scripts are available at https://github.com/Rowena-h/StarshipTrees.

## ACKNOWLEDGEMENTS

Many thanks to Javier Palma-Guerrero and Tania Chancellor for valuable discussion as part of the bilateral Earlham Institute–Rothamsted Research take-all working group. We also thank Neil Hall for his continued support and guidance.

